# Maternal Effects on Postembryonic Neuroblast Migration in *C. elegans*

**DOI:** 10.1101/2025.03.17.643740

**Authors:** Hoikiu Poon, Chaogu Zheng

**Affiliations:** School of Biological Sciences, The University of Hong Kong, Hong Kong SAR, China

## Abstract

Maternal effect genes mostly regulate early embryogenesis as their mRNAs or proteins are deposited into the oocytes to function during early embryonic development before the onset of zygotic transcription. Here, we report a case where a maternal effect gene regulates postembryonic neuroblast migration long after the early embryonic stages. We found that the defects of the Q neuroblast migration in *C. elegans* mannosyltransferase *dpy-19* mutants can be rescued by a maternal copy of the gene. Maternal *dpy-19* mRNAs are deposited into the oocytes and persist throughout embryonic development into the Q cells to regulate their migration in early larval stages. These mRNAs appeared to be remarkably stable, since long-term developmental arrest, changing the 3’UTR sequence, and mutations in genes involved in RNA binding and modification all had weak effects on the maternal rescue of the neuroblast migration defects. Since the defects can also be rescued by a zygotic copy of *dpy-19(+)*, our results suggest that postembryonic neurodevelopment is redundantly regulated by maternal and zygotic copies of the same gene.

## Introduction

Maternal effect refers to a phenomenon in which the phenotype of an organism is controlled not by its own genotype but by the genotype of its mother. Maternal effect genes are often essential for early developmental processes, since they code for RNAs and proteins important for early embryonic development before zygotic transcription takes place (WOLF AND WADE 2009; MITCHELL 2022). A subset of maternal effect genes does extend their influence into postembryonic development by controlling germline development. For example, maternally deposited *oskar* mRNAs in Drosophila eggs (LEHMANN 2016) and *bucky ball* mRNAs in Zebrafish oocyte (BONTEMS *et al*. 2009) help establish the germ plasm which is critical for germ cell specification. In *C. elegans*, maternal *pie-1* prevents somatic differentiation in early embryonic germline cells by repressing transcription (SEYDOUX *et al*. 1996) and the maternal *mes-2/3/6* genes regulate H3K27 methylation and are essential for germ-line development (XU *et al*. 2001; BENDER *et al*. 2004). Loss of these genes in the mother results in sterility in the offspring. However, very few studies reported maternal effect genes that regulate somatic cell differentiation in postembryonic development.

One example is the *clk-1* gene in *C. elegans*, which codes for a hydroxylase involved in the biosynthesis of ubiquinone (coenzyme Q10). The *clk-1* mutants have slower developmental rate due to the lack of ubiquinone, but the homozygous mutants produced by a heterozygous mother had a normal growth rate because the persistence of maternally deposited CLK-1 proteins enabled the synthesis of sufficient amounts of ubiquinone during development (BURGESS *et al*. 2003). In mammals, maternal nutrition during gestation affects postnatal development (KUSIN *et al*. 1992; HOFFMAN *et al*. 2016) and placenta-specific insulin-like growth factor 2 (IGF2) expression modulates fetal growth and contributes to postnatal metabolic homeostasis (CONSTANCIA *et al*. 2002; LOPEZ-TELLO *et al*. 2023). While these examples are mostly related to metabolism, instances of maternal effect genes regulating neuronal differentiation in postembryonic development are rare.

In this study, we report one such case where a maternal effect gene regulates postembryonic neuroblast migration in *C. elegans*. Previous studies of Q lineage development established a model for neuroblast migration in *C. elegans*, as the Q neuroblasts migrate, divide, and give rise to three types of neurons, including the mechanosensory touch receptor neurons (TRNs), oxygen-sensing neurons, and two interneurons (MIDDELKOOP AND KORSWAGEN 2014) (Fig. 1A). *dpy-19*, which codes for a mannosyltransferase, is required for the initial polarization of the Q cells by mediating the C-mannosylation of MIG-21 (a homolog of semaphorin-5A) and UNC-5 (a homolog of netrin receptor); C-mannosylation is essential for the secretion of soluble MIG-21 (HONIGBERG AND KENYON 2000; BUETTNER *et al*. 2013). The loss of *dpy-19* led to the mispositioning of the neurons derived from the Q lineage, including the TRN subtype PVM neuron. Through some serendipities, we found that the *dpy-19* homozygous mutants derived from the heterozygous mothers showed normal positioning of Q lineage descendants, indicating normal Q cell migration. We found that the *dpy-19* mRNAs but not proteins were maternally deposited into the oocytes and were passed down to the Q cells during embryonic development. Moreover, these maternal *dpy-19* mRNA appeared to be remarkably stable, since long-term developmental arrest, swapping the 3’UTR with non-maternal effect genes, and mutations in RNA stabilizing genes had only minor effects on the maternal rescue of the neuroblast migration defects. Overall, our study demonstrates that postembryonic neuronal development is safeguarded by the persistence of maternally deposited mRNAs.

**Figure 1.**
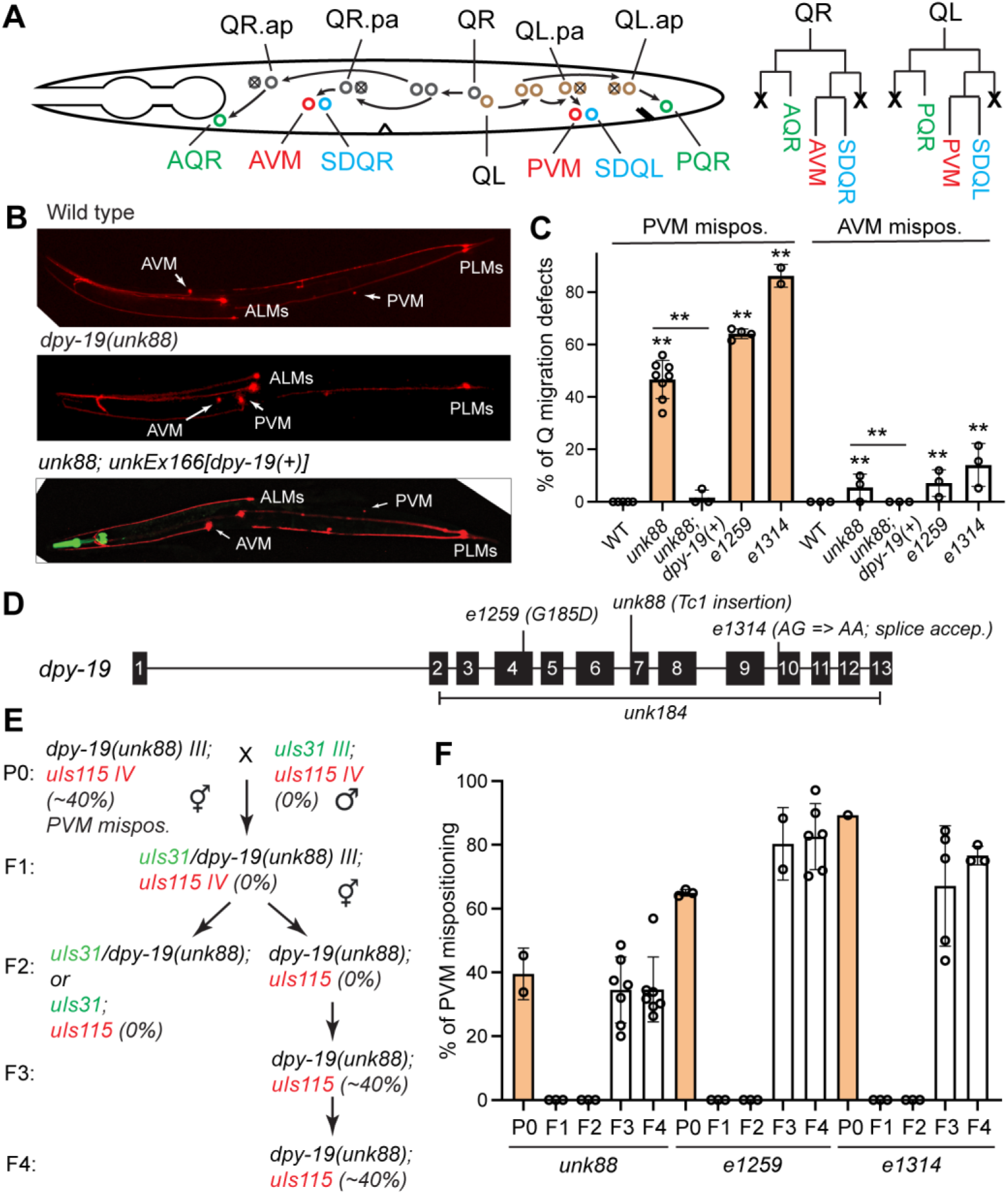
PVM positioning defects in *dpy-19* mutants are maternally rescued. (A) A schematic cartoon depicting the Q lineage and the migratory route of the Q neuroblasts and their descendants. (B) PVM is mispositioned to a more anterior position in *dpy-19(unk88)* mutants compared to the wild-type animals. This defect is rescued by an extrachromosomal array carrying a wild-type copy of *dpy-19*. TRNs are visualized using the reporter transgene *uIs115[mec-17p::TagRFP]*. (C) Percentage of animals showing PVM and AVM mispositioning phenotype as a result of QL and QR migration defects, respectively. (D) Gene structure of *dpy-19* and the molecular lesion caused by various alleles. (E) A cross scheme for maternal rescue experiment. *uIs31[mec-17p::GFP]* is integrated on chrIII and used as a marker for chrIII not carrying the *dpy-19* mutation. (F) Percentage of animals showing PVM mispositioning in each generation according to the labeling in (E) for various *dpy-19* alleles.

## Results

### A transposon insertion in *dpy-19* led to Q cell migration defects

In the process of studying the genetic control of TRN development, we analyzed the *ham-2(n1332)* allele, which was previously reported to affect HSN development (DESAI *et al*. 1988). We crossed the MT3149 *ham-2(n1332)* strain with the TRN marker *uIs115[mec-17p::TagRFP]* and segregated an autosomal allele that caused the mispositioning of the PVM neuron, which is a postembryonic TRN subtype derived from the QL cell lineage (Fig. 1A). In the wild-type animals, the PVM cell body was in the posterior half of the body, whereas in the mutants, ∼40% of the PVM was anteriorly displaced to a position close to the AVM cell body (Fig. 1B and C). This result was unexpected because *ham-2* is an X-linked gene, but the PVM phenotype was not X-linked. Using a combination of SNP mapping (DAVIS *et al*. 2005) and whole-genome resequencing, we mapped the mutation that caused PVM mispositioning to *dpy-19* and found a Tc1 insertion at the beginning of exon 7 (Fig. 1D and S1). The Tc1 transposon was found to be excised at a low frequency, which generates small indels (typically a 4-nucleotide insertion) in exon 7 (Fig. S1A). Interestingly, Tc1 was removed from the pre-mRNA likely due to the use of a cryptic splice site, which caused a frameshift deletion of 49 bp in exon 7 (Fig. S1B). We named the *dpy-19* allele *unk88*.

Tracing back to its origin, we found that the *ham-2(n1332)* allele was reported as a transposon insertion allele generated in the *mut-2* background (DESAI *et al*. 1988). One of the popular *mut-2* strains used in mutagenesis studies was MT3126 *mut-2(r459); dpy-19(n1347)* (RUSHFORTH et al. 1993). We genotyped *dpy-19* in MT3126 and found a Tc1 insertion at the same site as *unk88*, suggesting that *unk88* was likely to be *n1347*. Nevertheless, to avoid confusion, we still use *unk88* to refer to the *dpy-19* allele isolated from MT3149.

Previous results suggest that both QL and QR lacked clear polarization in *dpy-19* mutants (HONIGBERG AND KENYON 2000), suggesting that the positioning of the descendants of both QL and QR lineages may be similarly affected. We found that *dpy-19(unk88)* mutants mostly caused anterior displacement of the PVM cell body but rarely affected AVM positioning, suggesting that the QL migration may be more strongly disrupted by mutations in *dpy-19* than QR. To confirm that *dpy-19(unk88)* was indeed the phenotype-causing mutation, we injected a DNA construct containing the genomic fragment of *dpy-19(+)* into the mutants and found that the PVM mispositioning phenotype was fully rescued in the transformants (Fig. 1B and C).

### Maternal rescue of the Q migration defects in *dpy-19* mutants

In the process of analyzing the *unk88* allele, we noticed a maternal rescue of the PVM phenotype in the *dpy-19* mutants, where homozygous *dpy-19(unk88)* animals (F2) produced by the *dpy-19(unk88)/+* heterozygous mothers (F1) did not show the PVM mispositioning. We only started to observe the PVM phenotype in the homozygous F3 generation (Fig. 1E). Interestingly, *dpy-19(unk88)/+* heterozygotes produced by crossing homozygous hermaphrodites (P0) and wild-type males did not show the mispositioning of PVM cell body, suggesting that the phenotype was also rescued by the zygotic copy of *dpy-19(+)*. Moreover, *dpy-19(unk88)* animals produced by crossing *dpy-19* homozygous hermaphrodites with *dpy-19/+* heterozygous males showed the PVM phenotype, indicating the lack of paternal rescue (Fig. S2A). Since the PVM mispositioning phenotype had incomplete penetrance (∼40%) in *dpy-19* mutants, we also assessed whether the maternal phenotype of homozygous mothers (F3 generation) had an influence on the penetrance of the offspring and found no clear difference between the progeny of *dpy-19* mothers with and without the PVM phenotype (Fig. S2B-C).

Next, we compared *unk88* with two reference *dpy-19* alleles, *e1259* and *e1314*, for their phenotypes on Q neuroblast migration. *e1259*(G185D) is a missense allele in exon 4, and *e1314* is a splice acceptor mutation that affects the splicing of exon 10 (Fig. 1D). Both *e1259* and *e1314* alleles showed strong defects in QL migration, indicated by the anterior displacement of the PVM cell body in 60-90% of the animals (Fig. 1C). About 10-20% of AVM cell body was mispositioned to the posterior half of the animal in the mutants, suggesting that QR migration was also affected, although to a lesser extent. We also observed the maternal rescue of the PVM positioning defects in *e1259* and *e1314* mutants, since F2 homozygotes from the F1 *dpy-19/+* heterozygotes did not show any PVM phenotype (Fig. 1F).

*dpy-19* was originally identified by Brenner (1974) as a gene that, when mutated, causes the animals to be short and dumpy. Indeed, *e1259* and *e1314* mutants resulted in a temperature-dependent dumpy phenotype; when animals were grown at 20°C or above, their lengths were significantly lower than the wild type (Fig. S3A and B). Importantly, the dumpy phenotype of *dpy-19* mutants was also maternally rescued (Fig. S3C). Nevertheless, we did not observe an obvious dumpy phenotype in *unk88* mutants, suggesting that *unk88* may be a weaker allele than *e1258* and *e1314*. RT-PCR results showed that the *dpy-19* transcripts levels were comparable among the wild-type, the *unk88* mutants, and the *e1259* missense mutants, but were reduced in the *e1314* splice acceptor mutants (Fig. S3D).

In addition to AVM and PVM, the Q lineages also give rise to four other neurons: the oxygen-sensing neuron AQR and PQR, and the interneurons SDQL and SDQR. Defects in Q cell migration would also lead to the mispositioning of these neurons. Indeed, we observed the anterior displacement of PQR and SDQL cell bodies in ∼40% of the *dpy-19(unk88)* mutants and the posterior displacement of AQR and SDQR in 5∼10% of the mutants (Fig. 2A and B). As expected, the QL descendants (PQR, SDQL, and PVM) were often displaced together in the same animal, and the QR descendants (AQR, SDQR, and AVM) were also displaced together, confirming that mutations in *dpy-19* affected the initial migration of the Q neuroblasts. We also confirmed the maternal rescue of *dpy-19* mutants on the PQR and SDQL mispositioning phenotype, suggesting that the postembryonic development of the entire Q lineage is regulated by maternal DPY-19 (Fig. 2B).

**Figure 2.**
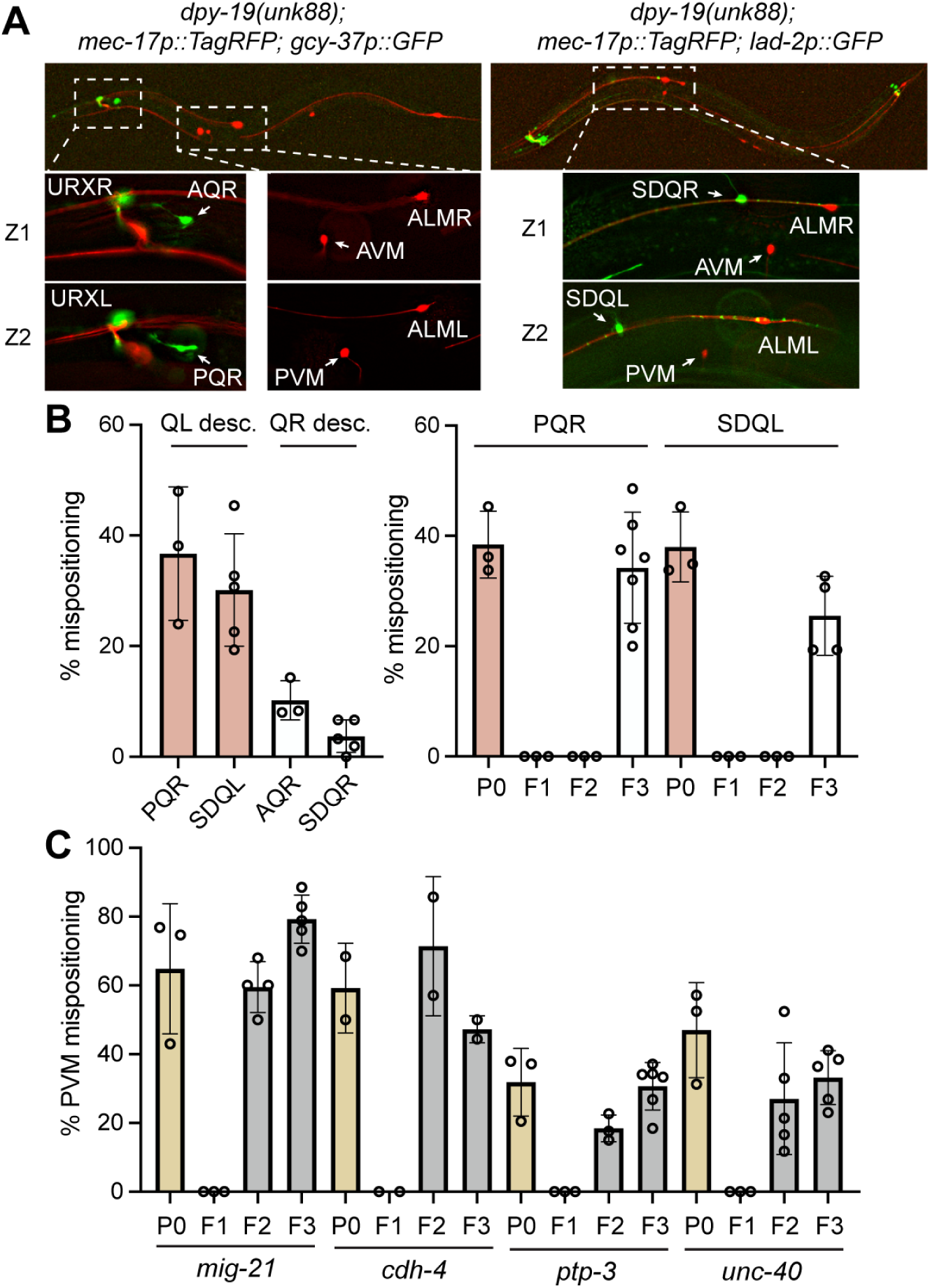
Maternal effects of *dpy-19* affect the positioning of all Q lineage descendants. (A) The positions of QL descendants (PVM, PQR, and SDQL) were all affected by *dpy-19(unk88)* mutation in the same animal. PQR, labeled by *iaIs25[gcy-37p::GFP]*, was normally positioned around the tail and was mispositioned to the head in *dpy-19* mutants; SDQL, labeled by *uIs130[lad-2p::GFP]*, was normally positioned in the posterior half of the animal and was mispositioned to the anterior in *dpy-19* mutants. Z1 and Z2 showed two focal planes of the same animal. (B) Percentage of *dpy-19(unk88)* animals showing PQR, SDQL, AQR, and SDQR cell body mispositioning phenotypes. Percentage of animals showing PQR and SDQL mispositioning in each generation in a cross similar to the one in Figure 1E. P0 and F3 animals were *dpy-19(unk88; M-)*, whereas F2 animals were *dpy-19(unk88; M+)*. (C) Test for maternal effects of four other genes involved in QL migration using a cross similar to the one in Figure 1E. *mig-21(u787)*, *cdh-4(hd40)*, *ptp-3(mu256)*, and *unc-40(n324)* alleles were used.

The C-mannosyltransferase DPY-19 promotes Q cell polarization by promoting the mannosylation and solubilization of the thrombospondin type-1 repeat (TSR)-containing membrane protein MIG-21 (BUETTNER *et al*. 2013). Mutations in *mig-21* caused a similar PVM displacement as *dpy-19* mutants (MIDDELKOOP *et al*. 2012), but the PVM phenotype in *mig-21* mutants were not maternally rescued (Fig. 2C). Moreover, UNC-40/DCC receptor, PTP-3/LAR-like receptor tyrosine phosphatase, and CDH-4/Cadherin also function in the same pathway as MIG-21 in regulating Q polarization (SUNDARARAJAN AND LUNDQUIST 2012; SUNDARARAJAN *et al*. 2014). Although their mutants all showed PVM mispositioning, none of them had maternal rescues (Fig. 2C). So far, to our knowledge, *dpy-19* appears to be the only Q cell regulating gene that displays maternal effects.

### *dpy-19* mRNAs are maternally deposited into embryos

Maternal effects are generally mediated by maternal deposits of proteins or mRNAs or epigenetic modifications, such as DNA methylation and histone modifications (MITCHELL 2022). To explore these mechanisms, we first examined a strain with GFP knock-in at the endogenous *dpy-19* locus (EBBING *et al*. 2019) and found no DPY-19::GFP signal in the early embryos, suggesting that there is no maternal deposition of DPY-19 proteins (Fig. 3A).

**Figure 3.**
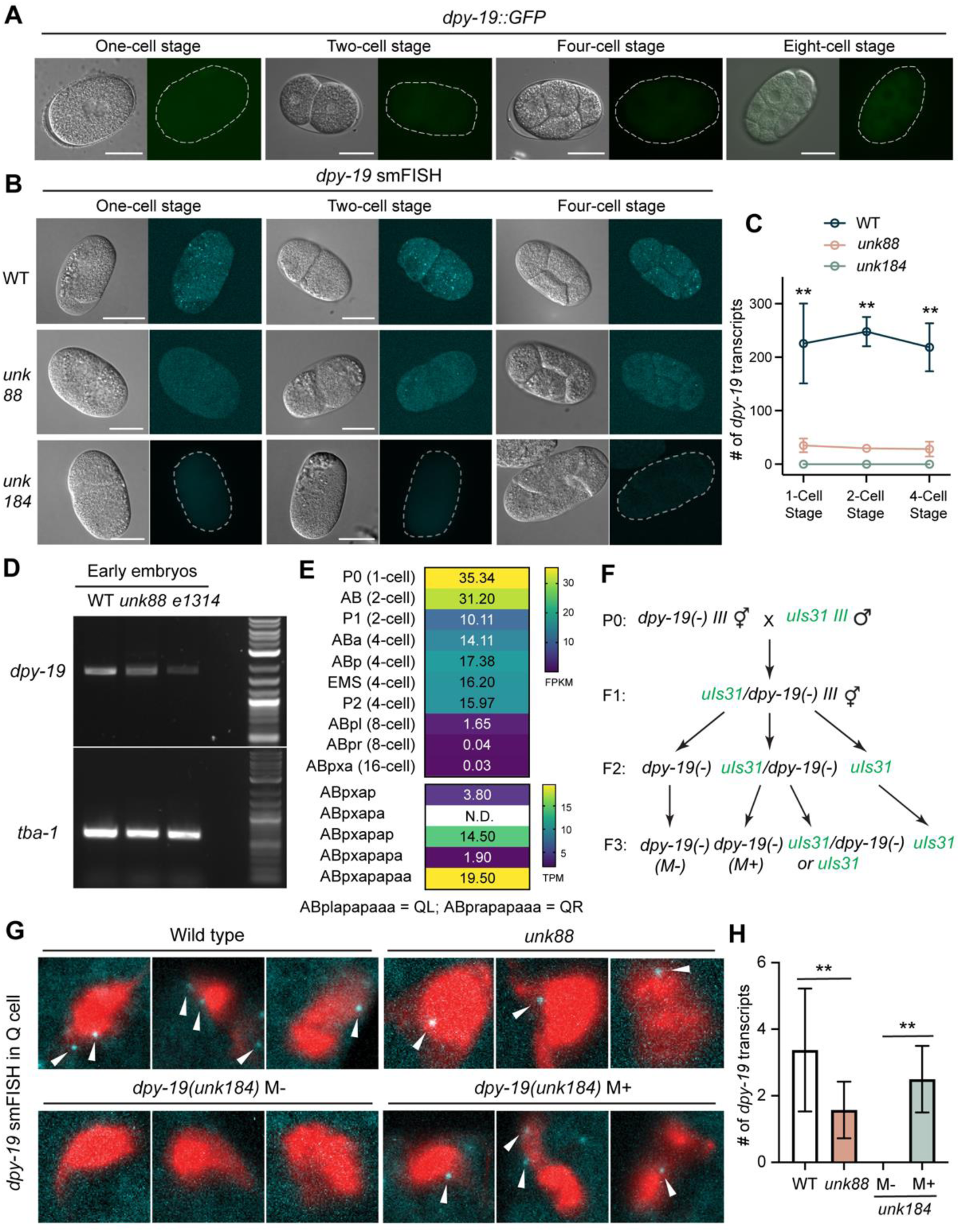
Maternally deposited *dpy-19* mRNAs persisted throughout development. (A) DPY-19::GFP expressed from an endogenous GFP knock-in allele *hu257[dpy-19::gfp::SEC::3xflag]* did not show any GFP signal in early embryos. (B) smFISH staining against *dpy-19* in wild-type, *unk88*, and *unk184* (the *dpy-19* deletion allele; Figure 1D) animals at early embryonic stages before the onset of zygotic transcription. Scale bars = 20 μm. (C) The number of all smFISH signals (i.e., fluorescence puncta) in each embryo counted through a Z stack of 10-15 pictures. Double asterisks indicate significant difference (*p* < 0.01) between the wild type animals and each of the two mutants in a post-ANOVA Dunnett’s test. (D) RT-PCR of *dpy-19* in the cDNA library of early embryos (unlaid eggs at 32-cell stage or earlier). (E) *dpy-19* expression levels extracted from published single-cell transcriptomic data of the embryos (TINTORI *et al*. 2016; PACKER *et al*. 2019). ABpl and ABpr lineages that give rise to QL and QR, respectively, are grouped into the same embryonic cell types as ABpx. (F) The cross scheme to examine the maternal *dpy-19* mRNAs using smFISH; *dpy-19(unk184)* was used as the *dpy-19(-)* knockout allele. We included the transgene *ayIs9[egl-17p::GFP]*, a Q cell marker, in the background to facilitate the identification of the Q cells. (G) smFISH against *dpy-19* in the Q cells of *dpy-19(-)* animals with or without maternal rescue according to the cross in (F). The wild-type and *unk88* animals served as controls; GFP signal from *ayIs9* was pseudo-colored in red to allow the visualization of smFISH signals (in cyan and indicated by arrow heads). (H) Quantification of the smFISH results from (G). Double asterisks indicate statistical significance (*p* < 0.01) in a post-ANOVA Tukey’s test.

To check for maternal deposition of *dpy-19* mRNAs, we performed single-molecule fluorescent *in situ* hybridization (smFISH) to quantitatively detect *dpy-19* mRNAs in early embryos before zygotic transcription occurs. For a negative control, we generated a *dpy-19* knockout mutant (*unk184*) through CRISPR/Cas9-mediated gene editing (Fig. 1D). The *unk184* allele was used as *dpy-19(-)* in the following studies. Using smFISH, we could clearly detect >200 *dpy-19* mRNAs in early embryos (i.e., zygote, 2-cell stage, and 4-cell stage embryos) in the wild-type animals, much reduced mRNAs levels (∼20 per embryo) in *unk88* animals, and virtually no mRNA signal in the *dpy-19* knockout mutants (Fig. 3B and C). We also conducted RT-PCR on the early embryos of up to the 8-cell stage and found *dpy-19* expression in the wild-type embryos; *unk88* and *e1314* mutants showed reduced embryonic mRNA levels (Fig. 3D). The above results indicate that *dpy-19* mRNAs are maternally deposited into the oocyte to mediate maternal effects.

Single-cell transcriptomic studies (TINTORI *et al*. 2016; PACKER *et al*. 2019) also confirmed *dpy-19* expression in all early embryonic cells from zygote to 4-cell stage, as well as most of the lineage precursors that give rise to the Q neuroblasts (Fig. 3E). These results at the single-cell level mapped out the lineage routes through which maternally deposited *dpy-19* mRNA could be passed to the Q neuroblasts of the offspring to produce DPY-19 proteins. To confirm this inheritance, we used smFISH to detect maternally deposited *dpy-19* mRNAs in the Q cells of *dpy-19(-)* animals produced by the *dpy-19(-)/+* heterozygous mothers and, indeed, found mRNA signals that overlap with the Q cells (QL and QR), labeled by *ayIs9[egl-17p::GFP]* (Fig. 3F-H). These results suggested that the absence of Q cell migration defects in the F2 *dpy-19(-)* homozygous animals was due to the deposition of wild-type *dpy-19* mRNA from the heterozygous mother (Fig. 3F-H).

In addition to maternal deposit, *dpy-19* is also likely to be transcribed zygotically, because the mRNA level of *dpy-19* is higher in mid- and late-stage embryos compared to the 1-cell embryo according to the modENCODE data (WormBase WS287). By studying embryonic transcriptomes, QUARATO *et al*. (2021) classified *dpy-19* as a “maternal stable” and “maternal and zygotic” gene, whose mRNAs level is detected in 1-cell embryos, is stably maintained in early embryos, and increases in late embryos. Zygotic expression of *dpy-19* is consistent with the zygotic rescue of PVM mispositioning phenotype in the cross progeny of *dpy-19* homozygous mothers and wild-type fathers (Fig. 1E).

Lastly, to explore whether DPY-19 may act through epigenetic modifications, we searched for potential DPY-19 substrates by scanning the *C. elegans* proteomes for the “WXXWXXW” motif, which is the glycosylation target recognized by DPY-19 (BUETTNER *et al*. 2013). The search yielded 41 possible DPY-19 targets, including MIG-21 and UNC-5, but none of them had direct connections with DNA methylation or histone modification (Table S1). Thus, we reason that DPY-19 is unlikely to produce maternal influence *via* epigenetic modifications.

### Maternal *dpy-19* mRNAs are likely stabilized *via* multiple mechanisms throughout development

The passage of maternally deposited *dpy-19* mRNA from oocyte to the Q cells suggests unusual stability of the mRNAs over a period of at least 14 hours (the time from fertilization to hatching, given that Q cells polarize and start to migrate shortly after hatching). We next attempted to identify the possible mechanisms that help stabilize *dpy-19* mRNA throughout embryogenesis. Inspired by a previous study (BURGESS *et al*. 2003), we first tested whether long-term developmental arrest at the L1 stage could destabilize maternal *dpy-19* mRNA and thus disallow the maternal rescue in *dpy-19(-)* mutants. Surprisingly, arresting the *dpy-19(-)* mutants derived from heterozygous mother at L1 stage for ≤ 30 days before recovery did not affect the maternal rescue at all, and we observed only ∼9% of the F2 mutants showing the PVM mispositioning phenotypes after 32 days of L1 arrest, compared to the ∼70% penetrance of the phenotype among the mutants derived from homozygous mothers (Fig. 4A). We reason that either *dpy-19* mRNA is incredibly stable, or the maternal *dpy-19* mRNA is only needed in the very early stage of L1 development before the starvation-induced arrest takes effect.

**Figure 4.**
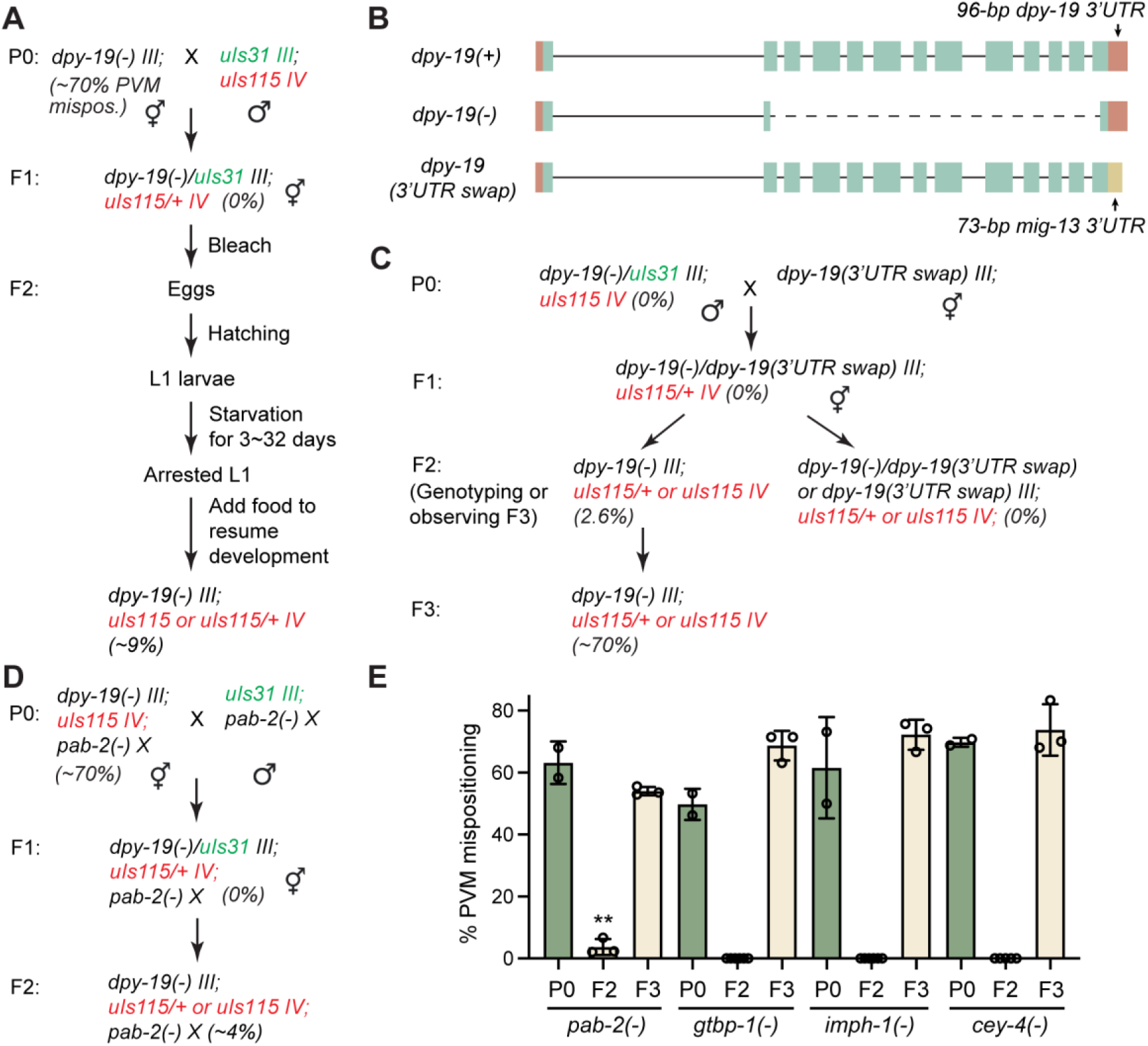
Maternal effect of *dpy-19* is robust against various perturbations. (A) An experimental scheme to test the effect of long-term developmental arrest on the maternal rescue of *dpy-19(unk184)* mutants. (B) Gene structure of *dpy-19(unk184)*, referred to as *dpy-19(-)*, and *dpy-19(syb8232)*, referred to as *dpy-19(mig-21 3’UTR)*, in which the *dpy-19* 3’UTR is replaced by *mig-21* 3’UTR at the endogenous locus. (C) A cross scheme to test the effects of the 3’UTR swap on the maternal rescue of the PVM mispositioning defects in *dpy-19(unk184)* mutants. (D) A representative scheme to test various genetic backgrounds for their impact on the *dpy-19* maternal rescue using *pab-2* as an example. (E) The percentage of animals with PVM mispositioning defects at different generations (P0 and F3 represented M-, while F2 represented M+) under *pab-2(ok1851)*, *gtbp-1(ax2029)*, *imph-1(tm1623)*, and *cey-4(ok858)* backgrounds using the cross schemes detailed in (D).

Since the 3’UTR often regulates the stability of mRNAs, including maternally deposited mRNAs (MISHIMA AND TOMARI 2016; DAI *et al*. 2019), we next tested whether *dpy-19* 3’UTR has similar effects. We swapped the 96-bp sequence encoding the *dpy-19* 3’UTR with a 73-bp fragment encoding the *mig-21* 3’UTR at the endogenous *dpy-19* locus through CRISPR/Cas9-mediated gene editing (Fig. 4B). The rationale was that the *mig-21* mutants caused similar PVM phenotypes but did not show any maternal effects. This swap did not affect *dpy-19* function, since we did not observe any phenotypes in the *dpy-19* mutants with *mig-21* 3’UTR. We then generated heterozygotes carrying one *dpy-19 (3’UTR swapped)* allele and one *dpy-19(-)* allele and found that among their *dpy-19(-)* homozygous mutant progeny, only 2.6% (n = 228) showed the PVM positioning defects, suggesting that the 3’UTR of the gene does not contribute significantly to the maternal rescue (Fig. 4C). Furthermore, the starvation-induced L1 arrest did not cause any stronger effects on the phenotype in the *dpy-19(-)* animals produced by the *dpy-19(-)/dpy-19(3’UTR swapped)* heterozygotes compared to the mutants derived from *dpy-19(-)/+* mothers (8% vs. 9%), supporting the idea that the *dpy-19* 3’UTR does not play a significant role in stabilizing its mRNA.

Lastly, we hypothesized that certain RNA-binding proteins may be involved in stabilizing the maternal *dpy-19* mRNA and preventing it from degradation during embryogenesis. We chose to test four genes, including *cey-4* (homolog of mammalian YBX1 and YBX2), *gtbp-1* (homolog of G3BP1), *imph-1* (homolog of IGF2BP), and *pab-2* (homolog of PABPC1), because of their functions in stabilizing target mRNA and interacting with m^6^A or m^5^C modified RNAs (BOO AND KIM 2020; NOMBELA *et al*. 2021). Although whether these RNA modifications exist on the mRNA of *C. elegans* is still under debate, recent studies found that m^6^A modification on the pre-mRNA of *sams-3* regulates mRNA splicing (MENDEL *et al*. 2021; WATABE *et al*. 2021). Thus, although these modifications may not be widely present in mRNAs, they may still occur on selected mRNA species. We first crossed the null alleles of the above genes with *dpy-19(-)* and confirmed that the PVM mispositioning phenotype was not affected in the double mutants. We then conducted the test for maternal rescue in the genetic background that is depleted of these RNA-binding proteins (Fig. 4D). We found that only the depletion of *pab-2* suppressed the maternal rescue in ∼4% (n = 152) of F2 *dpy-19(-)* mutants arising from heterozygous mothers, while the loss of the other three had no effects (Fig. 4E). *pab-2* codes for a homolog of PABPC1, which not only binds to the 3’ poly(A) tail of mRNAs (SACHS AND DAVIS 1989) but also interacts with m^6^A and m^5^C modified mRNAs through IGF2BP and YBX1, respectively (YANG *et al*. 2019; ZHANG *et al*. 2021). The fact that the loss of *imph-1/IGF2BP* and *cey-4/YBX1* did not produce similar effects suggests that the two RNA modifications may not be involved in stabilizing the *dpy-19* mRNAs, or each alone is not sufficient. The weak effects of *pab-2* may also be due to the redundancy with *pab-1*, which is an essential gene.

Given that the various treatment and genetic perturbations above did not significantly disrupt the maternal rescue of *dpy-19(-)* mutants, we reason that the maternal *dpy-19* mRNA may be stabilized by multiple and potentially redundant mechanisms or some unknown factors.

## Discussion

By tracking the Q lineage descendants in *C. elegans*, we showed that neuroblast migration in postembryonic neurodevelopment is regulated by maternally deposited *dpy-19* mRNAs. These mRNAs are so stable that they can be passed down from the zygote all the way to the embryonically born Q neuroblasts through ten rounds of cell divisions that take ∼720 minutes. These mRNAs then persist throughout the rest of embryonic development and the first few hours of larval development to regulate the initial polarization of the Q cells. Thus, the maternal *dpy-19* mRNAs can carry out their functions 15-16 hours after the cleavage of the zygote, which suggests remarkable perdurance given that the life cycle of the animal is merely 3 days. Although we found that the 3’UTR of *dpy-19* and the poly(A)-binding protein PAB-2 contribute to *dpy-19* mRNA stability, their contributions were relatively minor. It remains unclear what mechanisms allow *dpy-19* mRNA to survive the maternal mRNA clearance and persist throughout development. Interestingly, a zygotic copy of *dpy-19(+)* can also rescue the Q neuroblast migration defects in *dpy-19* mutants, suggesting that the maternal *dpy-19* mRNA is sufficient but not necessary for Q cell migration. This finding also suggests that the postembryonic neuronal development is a robust process redundantly regulated by both maternal and zygotic genes.

A few other examples of maternal genes regulating postembryonic development came from early screens for maternal-effect viable mutants in *C. elegans* (HEKIMI *et al*. 1995). For instance, the *clk* genes regulate embryonic and post-embryonic development, reproduction, and rhythmic behaviors, and their mutant phenotypes are fully rescued by maternal contributions (BENARD *et al*. 2001; BURGESS *et al*. 2003); *clk-1* codes for a 5-demethoxyubiquinone hydroxylase and *clk-2* codes for a telomere maintenance protein. Another example is the gene *mau-2*, which codes for a protein with predicted sister chromatid cohesin loading and dsDNA binding activities. *mau-2* mutants showed defects in larval development, locomotion, neuronal migration, and axonal guidance, which were all rescued by maternally deposited *mau-2* mRNAs (TAKAGI *et al*. 1997). MAU-2 regulates the positioning and axonal guidance of both embryonically and post-embryonically derived neurons including the TRN subtype AVM, and the AVM guidance defects in *mau-2* can also be rescued through a zygotic copy of the gene expressed cell-autonomously (BENARD *et al*. 2004). Given that 80 of the 302 neurons in the *C. elegans* adult hermaphrodite are derived post-embryonically, we suspect that the differentiation of many neurons in the nervous system may be subjected to both maternal and zygotic controls.

*dpy-19* codes for a conserved C-mannosyltransferase that attaches an α-mannose to the tryptophan residue in the TSR domain of the substrate (BUETTNER *et al*. 2013). C-mannosylation regulates protein folding and secretion (INAI *et al*. 2020). In the Q cells, DPY-19 glycosylates the TSRs in UNC-5/Netrin receptor and MIG-21/Sema5A for their stability and secretion (BUETTNER *et al*. 2013), which subsequently regulate Q cell polarization and migration. Interestingly, the DPY-19 function in neuronal migration appears to be conserved across species, as WATANABE *et al*. (2011) showed that the mouse homolog DPY19L1 is highly expressed in the developing glutamatergic neurons in the mouse embryonic cerebral cortex and regulates their radial migration. In humans, the deletion of DPY19L2 causes globozoospermia, a male infertility condition characterized by round-headed spermatozoa due to defective sperm head elongation and acrosome formation (HARBUZ *et al*. 2011; KOSCINSKI *et al*. 2011). Mechanistic studies in mice found that DPY-19L2 was located to the inner nuclear membrane of the spermatids and helped anchor acroplaxome to the nuclear envelope (PIERRE *et al*. 2012). Whether mammalian *dpy-19* homologs also show unusual mRNA stability and have maternal effects await further investigations.

## Materials and Methods

### Strains and transgenes

*C. elegans* wild-type (N2) and mutant strains were maintained as previously described (BRENNER 1974). Most of the experiments were performed at 20°C on NGM plates seeded with *E. coli* (OP50) as food source unless otherwise indicated. Transgene *uIs115[mec-17p::TagRFP]* was used to visualize the TRNs and the transgene *uIs130[lad-2p::GFP]* was used to visualize SDQ neurons. *uIs31[mec-17p::GFP]*, which is integrated on chromosome III and labels the TRNs in green, was used as a chr III marker. *dpy-19(e1259)*, *dpy-19(e1314)*, *mig-21(u787)*, *cdh-4(hd40)*, *ptp-3(mu256), unc-40(n324), pab-2(ok1851),* and other mutants were obtained from the Caenorhabditis Genetics Center. A list of strains used in this study can be found in Table S2.

### Genomic mapping of *unk88*

We first used a set of chromosome markers to test their linkage with *unk88* and found that the allele was located on chr III. We then used a previously published single-nucleotide polymorphism (SNP) two-point mapping method (DAVIS *et al*. 2005) to narrow down the genomic location of *unk88* by crossing the mutant strain with the genetically divergent CB4856 strain. This approach allowed us to map *unk88* between −19 and 4 on chr III. We then outcrossed the strain against the wild-type animals carrying *uIs115* and conducted whole-genome resequencing of the outcrossed strain and identified a 4-nt insertion (III: 8661495 GTA => GTACATA) in *dpy-19*. Genotyping of the *dpy-19* locus revealed the Tc1 transposon insertion and the 4-nt insertion appeared to have resulted from the self-excision of the transposon.

To rescue the *dpy-19(unk88)* mutant phenotype, we cloned a 11.7-kb genomic fragment including a 3-kb *dpy-19* promoter and the entire coding region and a 1-kb sequence downstream of the stop codon into pUC57. We then injected the *dpy-19(+)* construct into *dpy-19(unk88); uIs115* to obtain the transgenic animals with extrachromosomal array and examined the PVM positions.

### CRISPR/Cas9-mediated gene editing

To create the *dpy-19* deletion allele, we used CRISPR/Cas9-mediated gene editing to make cuts at two separate sites (exon 2 and exon 13) of the endogenous *dpy-19* locus. The two target sequences were 5’-GGTAGTTGATGTATCCAACG-3’ and 5’-ATGTGTTTATCGTGGAATGT-3’, respectively. The single guide RNAs (sgRNAs) were synthesized using the NEB EnGen sgRNA Synthesis Kit (E3322V) and were injected together with recombinant Cas9 (EnGen S. pyogenes Cas9 NLS from NEB, M0646T) into the *C. elegans*. We used pCFJ104 (*myo-3p::mCherry*) as the co-injection marker, and the transformants with red muscles were genotyped for any deletion in *dpy-19*. The allele that deleted the sequence between the two cut sites was named *unk184*.

To create the *dpy-19* (*mig-21 3’UTR*) allele, we chose the CRISPR/Cas9 target sequence 5’-AAATAGTTTATAATGCTAAT-3’ close to the stop codon of *dpy-19* and created sgRNA targeting the site and co-injected repair template that replaced the *dpy-19* 3’UTR (5’-tttttttctattttgttttt aaatttatttatttagttccagtatttttctgtaattccaaaacgatgaaatcaaatgaaccggtactgtatgttg-3’) with the *mig-21* 3’UTR (5’-atttcggatgctttcgagaacagtctctgtctgcccatttctcacgccacgataataaaagttatcattgatc-3’). The gene editing experiment was performed by SunyBiotech (Fuzhou, China) and the resulted allele was named *syb8232*. We verified the sequence through genotyping after receiving the strain.

### RT-PCR

Young embryos of N2, *unk88*, and *e1314* were collected in multiple batches. Animals were bleached to synchronize growth and grown on OP50-seeded 100mm NGM plates under 15°C to ensure abundant eggs were produced. When eggs were visible inside the gonad and ideally before gastrulation (and there were little to no eggs on the plate), worms were washed and bleached to collect the eggs. RNA extraction was performed using TRIzol reagent (Thermo Fisher) according to standard protocol. cDNAs from different samples were normalized using their *tba-1* expression pattern visualized on a gel. PCR against *dpy-19* was conducted using primers that bind to the exon 1-2 and exon 12-13 junctions. The PCR products were visualized on an agarose gel after electrophoresis and semi-quantitatively measured by comparison between the *dpy-19* mutants and the wild type. Primer information can be found in Table S2. Similar RT-PCR experiments were also conducted on mixed stage samples.

### Single-molecule *in situ* hybridization (smFISH)

To visualize the endogenous *dpy-19* mRNAs, we designed the smFISH probes against the *dpy-19* transcript using the Stellaris RNA FISH Probe Designer from Biosearch Technologies (Petaluma, CA) and synthesized the probes with Quasar 670 dye. The staining was conducted according to the procedures detailed in “Stellaris RNA FISH Protocol for *C. elegans*” by Biosearch Technologies. Eggs were obtained from gravid adults of N2, *unk88* and *unk184* strains. Samples were fixed in 37% formaldehyde, stored in 70% ethanol overnight, hybridized with a probe-containing hybridization buffer overnight, and then washed and imaged using the Leica DMi8 Inverted Microscope. 1-cell, 2-cell, and 4-cell stage eggs were visualized and quantified for the number of RNA signals detected. Each egg was imaged using the Z-stack function and one image was captured every 1.5 µm. A total of 10-15 images were taken for each sample. As each fluorescent puncta represents individual RNA molecules, we counted all puncta from the stack of images to obtain the total number of *dpy-19* transcripts in each sample.

To observe the presence of maternal *dpy-19* mRNAs in the Q lineage, we first conducted smFISH against *dpy-19* in a strain carrying a Q cell marker *ayIs9[egl-17p::GFP]* and the colocalization of the smFISH signal with GFP served as a positive control to show that *dpy-19* is indeed expressed in the Q cell. We then grew many *dpy-19(unk184)/uIs31 III* heterozygous animals to obtain F2 animals, which could be either non-fluorescent *dpy-19(unk184)* or fluorescent *dpy-19(unk184)/uIs31* and *uIs31*. We then bleached the F2 animals to get F3 eggs, which were grown to L1 and subjected to smFISH staining for *dpy-19*. The F3 animals that are derived from the non-fluorescent F2 *dpy-19(unk184)* animals did not inherit maternal *dpy-19* mRNA (i.e., M-), and the ones from F2 *dpy-19(unk184)/uIs31* animals inherited maternal dpy-19 mRNA (i.e., M+). In fact, in the non-fluorescent F3 animals, any *dpy-19* transcripts in the Q cells should be derived from maternally deposited *dpy-19* mRNAs.

### Tests of Maternal Rescue under various conditions

A typical maternal cross was carried out between *dpy-19* mutant hermaphrodites and wild-type males carrying the *uIs31[mec-17p::GFP]* transgene, which is located on the same chromosome as *dpy-19*. The green F1 heterozygous *dpy-19/uIs31* mutants were allowed to self-fertilize and produce non-green F2 *dpy-19* homozygotes, which were analyzed for PVM positioning. Both P0 parents carried *uIs115[mec-17p::TagRFP]* for the visualization of TRNs. Some F2 *dpy-19* homozygotes were picked onto fresh plates to produce F3 homozygotes, which were also examined. In the starvation experiment (BURGESS *et al*. 2003), we bleached the F1 animals to obtain F2 eggs, which were hatched in M9 and arrested at L1 stage on NGM plates, containing no food but ampicillin (100 µg/mL), kanamycin (50 µg/mL), and streptomycin (100 µg/mL) to prevent contamination for 3, 6, 9, 12, 15, 18, 30 and 32 days at 15°C. On the day of experiment, we flooded the plates with OP50-cotaining M9 to allow a thin layer of bacteria to form evenly, on which surviving animals fed until they reached day 1 adults. We examined the F2 *dpy-19* homozygous animals without green fluorescence for PVM positioning defects.

To test the effects of *dpy-19* 3’UTR, we first crossed *dpy-19(mig-21 3’UTR)* with *dpy-19(unk184)* to obtain the heterozygotes and then examined the *dpy-19(unk184)* homozygotes (by genotyping or observing dumpy animals in the F3) for potential PVM mispositioning phenotypes. To detect the effects of mutating genes that may regulate mRNA stability, we first crossed *pab-2(ok1851)* and *gtbp-1(ax2029)* with *dpy-19(unk184)* to create the double mutants. Since *cey-4* and *imph-1* are located on the same chromosome as *dpy-19*, we generated double mutants by deleting *dpy-19* in *cey-4(ok858)* and *imph-1(tm1623)* mutants using the same gene editing method described above. We then crossed the double mutants with the corresponding single mutants (i.e., *pab-2*, *gtbp-1*, *cey-4*, and *imph-1*, respectively) and analyzed the F2 progeny for potential disruption of maternal rescue.

### Quantification of PVM mispositioning

The AQR, AVM, and SDQR neurons in the wild-type animals reside in the anterior half of the worm’s body, while PQR, PVM and SDQL reside in the posterior half. “Anterior” is defined as any point anterior to the vulva, while “posterior” is defined as posterior to the vulva. In this study, “mispositioning” of a neuron means that an anterior neuron was observed in the posterior half of the body, or a posterior neuron was observed in an anterior half. We counted the number of animals showing mispositioned neurons in a population to obtain the penetrance of a mutant phenotype. Microscopic imaging was done on a Leica DMi8 inverted microscope equipped with a with a Leica K5 monochrome camera, and images were analyzed using the Leica Application Suite X software. In general, QL migration was more severely affected than that of QR in *dpy-19* mutant animals. So, we focused on the mispositioning of PVM, PQR, and SDQL neurons. To compare the penetrance across strains, we calculated the average percentage of animals showing the PVM mispositioning phenotype across at least three biological replicates and used a one-way ANOVA followed by either Dunnett’s test or Tukey’s honestly significant difference (HSD) test to find out statistical significance in multiple comparison.

## Acknowledgement

We thank Prof. Hendrik Korswagen at the Hubrecht Institute for providing the *dpy-19::GFP* strain and Prof. Erik Lundquist at the University of Kansas for providing the Q cell marker *ayIs9[egl-17p::GFP]* strain. This study is supported by funds from the National Natural Science Foundation of China (Excellent Young Scientists Fund for Hong Kong and Macau, 32122002), the Health Bureau of Hong Kong (HMRF 09201426 to C.Z.), and the Research Grant Council of Hong Kong (GRF 17105523, GRF 17106322, GRF 17107021, and CRF C7026-20G). Some strains used in this study were provided by the Caenorhabditis Genetics Center, which is funded by the NIH Office of Research Infrastructure Programs (P40 OD010440).

## AUTHOR CONTRIBUTIONS

C.Z. and H.P. conceived the study and wrote the manuscript. H.P. carried out most of the experiments. C.Z. did the initial characterization of the phenotype, secured funding, and supervised the project. Both authors read and approved the manuscript.

## DECLARATION OF INTERESTS

The authors declare no competing interests.

